# Bridging the gap between theory and data: the Red Queen Hypothesis for sex

**DOI:** 10.1101/637413

**Authors:** Sang Woo Park, Benjamin M Bolker

## Abstract

Sexual reproduction persists in nature despite its large cost. The Red Queen Hypothesis postulates that parasite pressure maintains sexual reproduction in the host population by selecting for the ability to produce rare genotypes that are resistant to infection. Mathematical models have been used to lay theoretical foundations for the hypothesis; empirical studies have confirmed these predictions. For example, Lively used a simple host-parasite model to predict that the frequency of sexual hosts should be positively correlated with the prevalence of infection. Lively *et al.* later confirmed the prediction through numerous field studies of snail-trematode systems in New Zealand. In this study, we fit a simple metapopulation host-parasite coevolution model to three data sets, each representing a different snail-trematode system, by matching the observed prevalence of sexual reproduction and trematode infection among hosts. Using the estimated parameters, we perform a power analysis to test the feasibility of observing the positive correlation predicted by Lively. We discuss anomalies in the data that are poorly explained by the model and provide practical guidance to both modelers and empiricists. Overall, our study suggests that a simple Red Queen model can only partially explain the observed relationships between parasite infection and the maintenance of sexual reproduction.

## 1 Introduction

Despite being the dominant mode of reproduction in multicellular organisms (Vrijenhoek, 1998; Whitton et al., 2008; Otto, 2009), sexual reproduction entails numerous costs (Lehtonen et al., 2012). The most commonly mentioned is the cost of producing males (Smith, 1978): as males cannot produce offspring, asexual lineages are expected to outgrow their sexual counterparts. This *two-fold cost of sex* (Smith, 1978) relies on the assumption that everything else is equal. Then, what drives the observed prevalence of sexual reproduction?

One explanation for the persistence of sexual reproduction is the Red Queen Hypothesis (RQH) (Bell, 1982). The RQH predicts that sexually reproducing hosts overcome the cost of sex under strong parasitic pressure by producing genetically diverse offspring that can escape infection (Haldane, 1949; Jaenike, 1978; Hamilton, 1980; Hamilton et al., 1990). Limited genetic diversity of asexually reproducing hosts allows for sexual reproduction to be maintained in the host population (Ashby and King, 2015).

Much of the theoretical literature has focused on determining qualitative conditions under which parasite selection can maintain sexual reproduction in the host population. Here, we describe a few important criteria. First, hosts and parasites must coevolve (Bell, 1982). Host-parasite coevolution creates a time-lagged selective advantage for rare host genotypes, creating an oscillation in genotypic frequencies (Clarke, 1976; Jaenike, 1978; Hamilton, 1980; Agrawal and Lively, 2001); this process promotes host genetic diversity and not necessarily sex (King et al., 2009; Dagan et al., 2013a; Ashby and King, 2015). Second, parasites must be highly virulent (May and Anderson, 1983). Although sexual and asexual hosts can coexist at intermediate virulence, sexual reproduction is unable to provide enough advantage to overcome the cost of sex against avirulent parasites (Howard et al., 1994). Finally, sexual hosts must be genetically more diverse than asexual hosts, as high clonal diversity may diminish the advantage of sexual reproduction (Lively and Howard, 1994; Lively, 2010c; Ashby and King, 2015).

Some theoretical studies have departed from the classical population genetics framework to study the effects of ecological and epidemiological context on the Red Queen dynamics. A few studies have suggested that incorporating more realistic ecological and epidemiological details may assist in supporting sexual reproduction in the host population (Lively, 2009, 2010b) and maintaining co-evolutionary cycles (Ashby and Gupta, 2014). In contrast, MacPherson and Otto (2018) showed that Red Queen dynamics (i.e., cycles in allele frequencies) fail to persist when an explicit epidemiological structure is taken into account with coevolutionary dynamics. Given the wide range of assumptions underlying these eco-evolutionary models, it may be difficult to understand how different ecological details of these models affect the overall maintenance of sexual reproduction; however, it is clear is that these details play important roles in shaping coevolutionary dynamics (Song et al., 2015; Haafke et al., 2016; Ashby et al., 2019).

Many empirical studies have focused on confirming predictions that stem from the RQH. Typical among them are rapid parasite evolution (Rauch et al., 2006), local adaptation (Lively, 1989; Morran et al., 2014; King et al., 2011; Gibson et al., 2016), time-lagged selection (Buckling and Rainey, 2002; Decaestecker et al., 2007; Koskella and Lively, 2009; Thrall et al., 2012; Koskella, 2013, 2014), and association between parasite prevalence and host reproductive mode (Lively, 1992; Vergara et al., 2013; Verhoeven and Biere, 2013). A key example is the snail population in New Zealand that serves as intermediate hosts for trematodes (Winterbourn, 1974; McArthur and Featherston, 1976). Through decades of work, Lively *et al.* have demonstrated that the population satisfies necessary conditions for the host-parasite coevolutionary dynamics and provides support for the hypothesis (e.g., Lively (1987, 1989); Dybdahl and Lively (1995b, 1998); Jokela et al. (2009); Vergara et al. (2014b); Gibson et al. (2016)). While field studies often provide only indirect evidence for the hypothesis, experimental systems can be used to directly test the hypothesis (Auld et al., 2016; Slowinski et al., 2016; Lynch et al., 2018; Zilio et al., 2018).

Even though the Red Queen Hypothesis has gained both theoretical and empirical support, there still remains a gap between theory and data. Many theoretical models rely on simplifying assumptions that are not applicable to natural populations and make indirect connections to empirical studies. For example, none of the Red Queen models reviewed by Ashby and King (2015) make statistical connections to empirical data. It is unclear how well these models capture coevolutionary dynamics in natural systems.

Theoretical models can still be used to make *qualitative* predictions about nature. Lively (1992) initially postulated that infection prevalence should be positively correlated with the frequency of sexual hosts and later formalized the idea using a mathematical model (Lively, 2001). The prediction has since been confirmed in several empirical studies, most of which are based on the snail-trematode system (Lively and Jokela, 2002; Kumpulainen et al., 2004; King et al., 2011; Vergara et al., 2013; McKone et al., 2016; Gibson et al., 2016). Surprisingly, the predicted correlation was not observed in a different snail-trematode system (Heller and Farstey, 1990; Ben-Ami and Heller, 2005, 2007, 2008; Dagan et al., 2013a).

Here, we try to help bridge the gap between theory and data in understanding the role of parasites in maintaining sexual reproduction in host populations and make quantitative inference about empirical systems. We extend the model used by Lively (2010b) to account for demographic stochasticity and include a simple metapopulation structure. Then, we fit the model to data sets from three field studies (Dagan et al., 2013a; McKone et al., 2016; Vergara et al., 2014b) using Approximate Bayesian Computation (ABC) to estimate biologically relevant parameters. We assess model fits and discuss discrepancies between a theoretical model and the observed data. Using biologically realistic parameters, we test for power (probability of observing a significant effect) to detect a positive correlation between frequency of sexual reproduction and prevalence of infection Lively (2001). We provide guidance for studying the RQH and discuss underlying factors that drive the correlation.

## 2 Methods

### 2.1 Data

We analyze three observational data sets (Vergara et al., 2014b; McKone et al., 2016; Dagan et al., 2013a) that represent interactions between freshwater snails and sterilizing trematodes. The New Zealand snail populations (Vergara et al., 2014b; McKone et al., 2016) consist of *Potamopyrgus antipodarum*, which serves as an intermediate host for trematodes, among which *Microphallus* sp. is most commonly found (Winterbourn, 1974; Lively, 1987). The Israeli snail populations (Dagan et al., 2013a) consist of *Melanoides tuberculata*, which also serves as an intermediate host for trematodes, including *Centrocestus* sp. (Ben-Ami and Heller, 2005), *Philophthalmus* sp. (Ben-Ami, 2006), and *Transversotrema patialense* (Ben-Ami et al., 2005). Diploid sexual females and polyploid asexual females coexist in both the New Zealand (Phillips and Lambert, 1989; Wallace, 1992; Dybdahl and Lively, 1995a) and the Israeli (Samadi et al., 1999) snail populations.

We choose to analyze snail-trematode systems because they have been extensively studied under the context of the Red Queen Hypothesis. The snail-trematode systems in New Zealand exhibit positive correlation between prevalence of infection and frequency of sexual hosts (Lively and Jokela, 2002; King et al., 2011; Vergara et al., 2013; McKone et al., 2016; Gibson et al., 2016); local adaptation (Lively, 1989; Lively et al., 2004; King et al., 2011); negative frequency-dependent selection (Dybdahl and Lively, 1995b, 1998; Jokela et al., 2009; Koskella and Lively, 2009); and periodic selection for sexual reproduction (Vergara et al., 2014b; Gibson et al., 2018). These results are consistent with the host-parasite coevolutionary theory behind the Red Queen Hypothesis. Therefore, we expect a basic Red Queen model to be able to mimic the observed dynamics reasonably well.

We included the Israeli population because there has been a consistent lack of positive correlation between the prevalence of infection and frequency of sexual hosts (Heller and Farstey, 1990; Ben-Ami and Heller, 2005, 2007, 2008; Dagan et al., 2013a), which sharply contrasts the observations from the New Zealand population. Comparison of these two populations with a mathematical model may allow us to better understand the underlying cause of the difference.

The data sets include proportions of infected snails, proportions of sexual/asexual snails, and locations of sampling sites; the data set from Vergara et al. (2014b) also includes sampling year. Whereas Vergara et al. (2014b) report infection status of snails specific to *Microphallus* sp., McKone et al. (2016) and Dagan et al. (2013a) do not distinguish among different species. The data sets collected by Dagan et al. (2013a) and Vergara et al. (2014b) are obtained from their Dryad repositories (Dagan et al., 2013b; Vergara et al., 2014a). The data set collected by McKone et al. (2016) is extracted from their figure.

### 2.2 Model

We model obligately sexual hosts competing with obligately asexual hosts in a metapopulation by extending the model introduced by Lively (2010b). Our model is a discrete time susceptible-infected (SI) model with natural mortality and virulence (defined as a reduction in offspring production among infected hosts). Metapopulation structure is included to model unobserved dynamics among different habitats; equivalently, each population can be considered as a sampling site. A similar metapopulation model was developed by Lively (2018) in order to study the local adaptation of parasites.

We do not model the life history of snail-trematode interactions explicitly. Modeling interactions of multiple trematodes species, each of which goes through a different life cycle, with a single snail species is extremely complicated. Instead, testing a basic model against data will allow us to identify the difference between theory and data more clearly.

All hosts are diploids with two biallelic loci controlling resistance; parasites are assumed to be haploids. Let 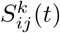 and 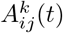 be the number of sexual and asexual hosts with genotype *ij* from subpopulation *k* at generation *t*, where the subscripts, *i* and *j*, represents host haplotypes: *i, j* ∈ {AB, Ab, aB, ab}. For simplicity, we drop the superscript representing the subpopulation and write *S*_*ij*_ (*t*) and *A*_*ij*_ (*t*); every population is governed by the same set of equations unless noted otherwise (e.g., when we account for the interaction between populations). Following Lively (2010b), the expected genotypic contribution (before recombination or outcrossing) by sexual hosts is given by

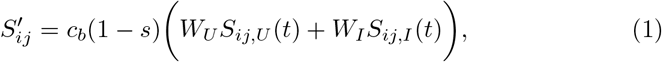

where *s* is the proportion of males produced by sexual hosts, and *S*_*ij,U*_ and *S*_*ij,I*_ are the number of uninfected and infected sexual hosts in a population. *W*_*U*_ and *W*_*I*_ represent their corresponding fitnesses where virulence is defined as *V* = 1 − *W*_*I*_ /*W*_*U*_. We allow for the cost of sex to vary by multiplying the growth rate by a scale parameter, *c*_*b*_, where 2/*c*_*b*_ corresponds to a two-fold cost of sex when *c*_*b*_ = 1 (Ashby and King, 2015). Recombination and outcrossing are modeled after incorporating genotypic contributions from other populations.

We define

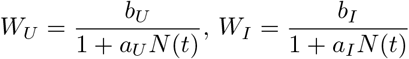

where *b*_*U*_ and *b*_*I*_ are the maximum number of offspring produced by uninfected and infected hosts, respectively, and *a*_*U*_ and *a*_*I*_ determine their corresponding strengths of density dependence (Smith and Slatkin, 1973; Lively, 2010b). For simplicity, we assume that *a*_*U*_ = *a*_*I*_ so that virulence can be defined strictly in terms of decrease in offspring production: *V* = 1 − *b*_*I*_/*b*_*U*_.

Asexual hosts are strictly clonal. The expected genotypic contribution by asexual hosts is given by

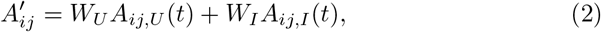

where *A*_*ij,U*_ and *A*_*ij,I*_ are the number of uninfected and infected asexual hosts in a population.

A proportion *ϵ*_mix_ of a population mixes with other populations. The expected number of offspring in the next generation (accounting for contributions from all populations) is given by

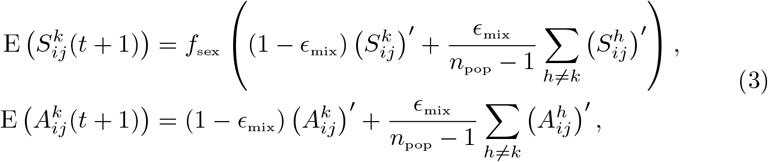

where *f*_sex_(*x*) is the function that models sexual reproduction, including recombination probability *r*_host_ and outcrossing, and *n*_pop_ is the number of populations modeled. The total number of sexual and asexual hosts in the next generation is given by a Poisson random variables with means specified previously. We also allow for stochastic migration in order to avoid fixation:

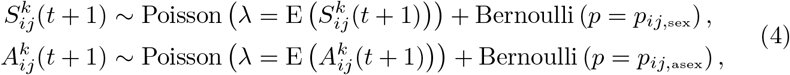

where *p*_*ij*,sex_ and *p*_*ij*,asex_ are the probabilities of a single sexual or asexual host with genotype *ij* entering a subpopulation.

Infection is modeled using the matching alleles model (Otto and Michalakis, 1998). We assume that snails are equally susceptible to parasites that match either haplotype. However, parasites must match the host haplotype at both loci in order to successfully infect a host. The expected number of infected hosts that are infected with parasite with genotype *i* at generation *t* is given by:

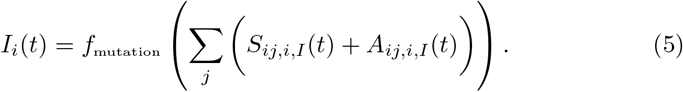

*S*_*ij,i,I*_(*t*) and *A*_*ij,i,I*_(*t*) represent the expected numbers of sexual and asexual hosts that have genotype *ij* and are infected with parasites that have genotype *i*. The function *f*_mutation_ models mutation at a single locus with probability *r*_parasite_ (Ashby and King, 2015). We further account for stochastic migration with probability *p*_*i*,parasite_ to avoid fixation:

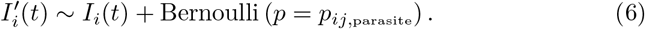

The total expected number of infectious contacts made by infected hosts within a population is given by 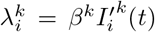, where *β*^*k*^ is the transmission rate of each population. We model mixing between subpopulations by allowing infected hosts to make contact with susceptible hosts in other populations; we assume that a proportion *ϵ*_mix_ of the infectious contacts are equally distributed among other subpopulations. The total amount of infectious contact, coming from hosts that carry genotype *i* parasite, that is received by susceptible hosts in population *k* is given by

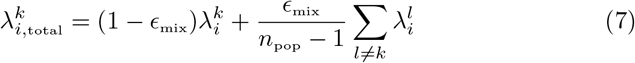

Then, the force of infection that a susceptible host with genotype *ij* experiences in generation *t* + 1 is given by

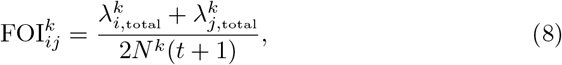

where 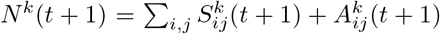 is the total number of hosts in generation *t* + 1. The probability that a susceptible host with genotype *ij* in population *k* becomes infected in the next generation is given by

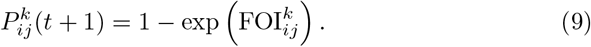

Finally, the number of infected hosts in the next generation is determined by a binomial process:

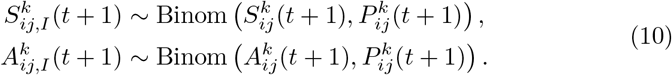

The expected number of infected hosts that have genotype *ij* and are infected by parasites with genotype *i* in the next generation is proportional to the amount of infectious contact that was made in the current generation:

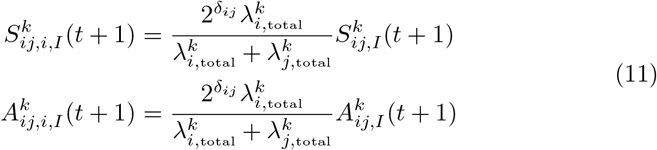

where a *δ*_*ij*_ is a Kronecker delta (*δ*_*ij*_ equals 1 when *i* = *j* and 0 otherwise).

### 2.3 Simulation design and parameterization

Many Red Queen models have focused on competition between a single asexual genotype and multiple sexual genotypes or have assumed equal genetic diversity between asexual and sexual hosts (see Ashby and King (2015) for a review of previous Red Queen models) but neither of these assumptions is realistic. In contrast, Ashby and King (2015) adopted a more realistic approach by incorporating stochastic external migration of an asexual genotype to a population and allowing for asexual genetic diversity to vary over time. Here, we combine these methods. We allow for stochastic external migration of asexual hosts with different genotypes into the system but fix the number of asexual genotypes (denoted by *G*_asex_) that can be present in the system. Since sexual hosts have limited genetic diversity in our model (diploid hosts with two biallelic loci yields a total of 10 genotypes), allowing for unlimited migration of asexual hosts will cause the sexual population to be easily outcompeted by the asexual population. Limiting asexual genetic diversity allows us to account for the intrinsic difference in sexual and asexual diversity and make a compromise between simple and realistic models. At the beginning of each simulation, a pool of *G*_asex_ asexual genotypes are sampled at random from the entire genotypic space; these are the genotypes that are available for external immigration into subpopulations. Then, we use our model-based comparison with data to estimate *G*_asex_ to test whether the data are consistent with the dynamics of a model that allows for more than a single asexual clone.

To account for differing number of sexual and asexual genotypes, we let

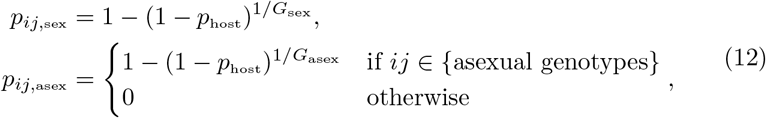

where *p*_host_ is the probability that at least one sexual or asexual host enters the population in a generation. We scale the probability of an infected host carrying parasite genotype *i* in a similar way for interpretability:

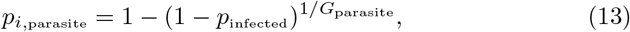

where *p*_infected_ is the probability that at least one infected host enters the population in a generation. The number of parasite genotypes *G*_parasite_ is equal to 4 (because parasites are assumed to be haploids with two loci).

Each simulation consists of 40 subpopulations. Every subpopulation is initialized with 2000 sexual hosts, of which 80 are infected. They are assumed to be in Hardy-Weinberg equilibrium where the allele frequency in each locus is equal to 0.5. In order to account for variation in infection rates among subpopulations, the transmission rate, *β*^*k*^, is randomly drawn for each subpopulation from a Gamma distribution with mean *β*_mean_ and coefficient of variation *β*_CV_. The simulation runs for 500 generations without the introduction of asexuals. At generation 501, 10 asexual hosts of a single genotype are introduced to each population (the asexual genotype introduced can vary across subpopulations) and simulation runs for a further 600 generations while allowing for stochastic migration of asexuals.

### 2.4 Approximate Bayesian Computation

We use Approximate Bayesian Computation (ABC) to estimate parameters from data (Toni et al., 2009). ABC relies on comparing summary statistics of observed data with those of simulated data; it is particularly useful when the exact likelihood function is not available. Each summary statistic (also referred to as a *probe*) represents an aspect of a system Kendall et al. (1999); probe matching can be more powerful than classical likelihood-based approaches because it allows to evaluate model fits based on biologically meaningful aspects of a system (Wood, 2010). We consider the mean proportion of infected and sexually reproducing snails in the system and variation in these proportions — measured by a coefficient of variation (CV) — across space (population) and time as our probes. As Dagan et al. (2013a) and McKone et al. (2016) only reported the proportion of males, the proportion of sexual hosts is assumed to be twice the proportion of males.

CV across space is calculated by first calculating mean proportions by averaging across time (generation) for each site (subpopulation) and then taking the the CV of these mean proportions. CV across time (generation) is calculated by first averaging proportions across space (subpopulation) at each generation and then taking the CV. For purely spatial data (Dagan et al. (2013a) and McKone et al. (2016)), CV across space is calculated without averaging across time.

Because ABC is a Bayesian method, we must specify prior distributions for all parameters. We use weakly informative priors for all estimated parameters except *c*_*b*_, a scale parameter for the cost of sex (see Table 1 for prior distributions used and parameters assumed). The prior distribution for the scale parameter is chosen so that 95% prior quantile range of cost of sex (2/*c*_*b*_) is approximately equal to the 95% confidence interval reported by Gibson et al. (2017). For brevity, all other parameters are assumed to be fixed. While it is a common practice to fix several parameters of a Red Queen model (e.g., Lively (2010b); Ashby and King (2015); Haafke et al. (2016); Ashby et al. (2019)), failing to explicitly account for uncertainty in all parameters of a model can lead to overly confident conclusions (Elderd et al., 2006).

**Table 1:**
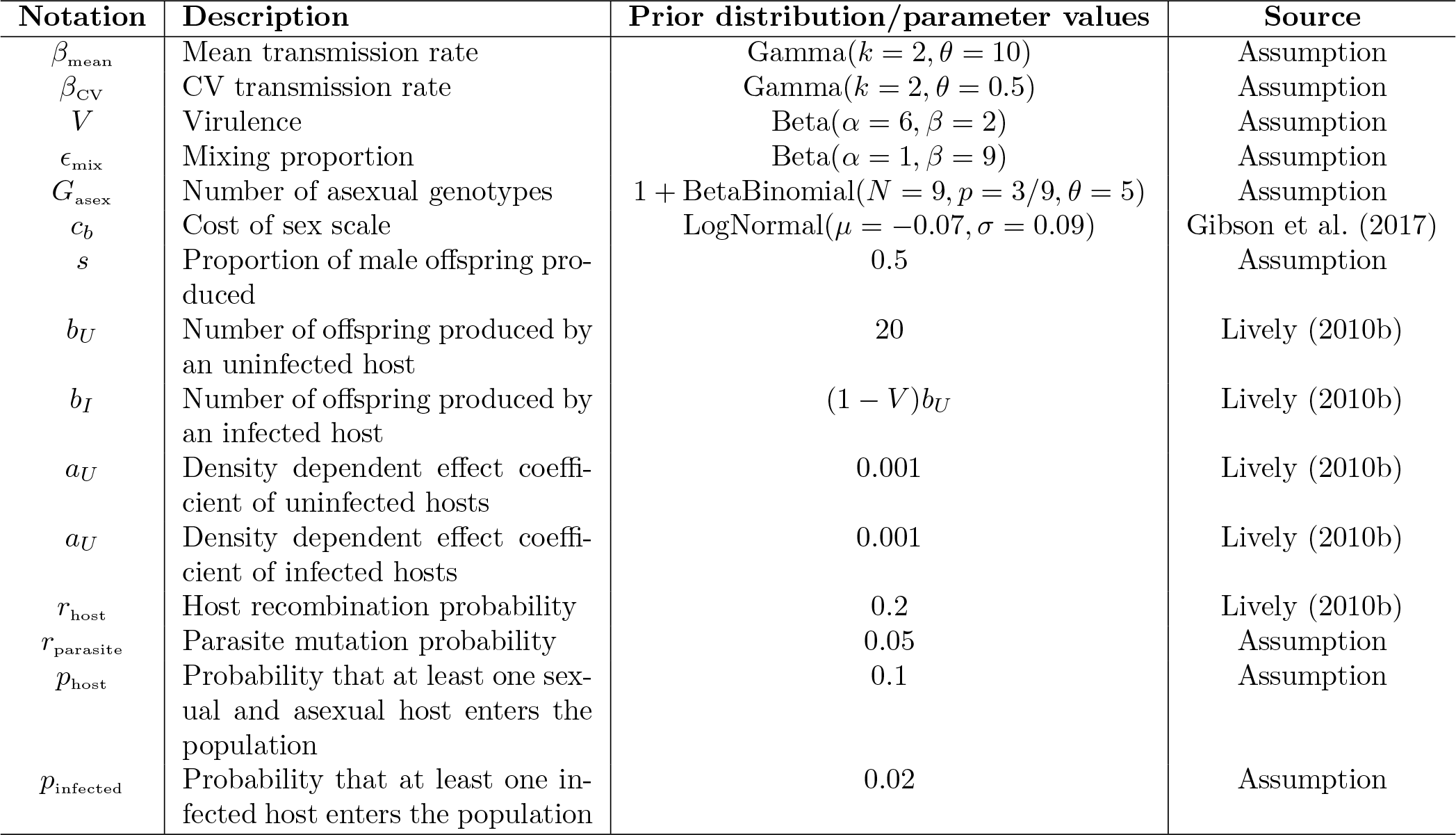
Parameter descriptions and values. Parameters with prior distributions are estimated via Approximate Bayesian Computation (ABC). Parameters *k* and *θ* in Gamma distribution represent shape and scale parameters where mean and squared CV are given by *kθ* and 1/*k*, respectively. *α* and *β* in Beta distribution represent shape parameters where mean and squared CV are given by *α*/(*α* + *β*) and *β/*(*α*^2^ + *αβ* + *α*). *N*, *p* and *θ* in Beta binomial distributions represent number of trials, probability of success, and overdispersion parameters (Morris et al., 1983). Parameters *μ* and *σ* in a log-normal distribution represent mean and standard deviation on a log scale. All other parameters are fixed throughout simulations.

| Notation | Description | Prior distribution/parameter values | Source |
| --- | --- | --- | --- |
| $\beta_{\text{mean}}$ | Mean transmission rate | Gamma($k = 2, \theta = 10$) | Assumption |
| $\beta_{\text{CV}}$ | CV transmission rate | Gamma($k = 2, \theta = 0.5$) | Assumption |
| $V$ | Virulence | Beta($\alpha = 6, \beta = 2$) | Assumption |
| $\epsilon_{\text{mix}}$ | Mixing proportion | Beta($\alpha = 1, \beta = 9$) | Assumption |
| $G_{\text{asex}}$ | Number of asexual genotypes | $1 + \text{BetaBinomial}(N = 9, p = 3/9, \theta = 5)$ | Assumption |
| $c_b$ | Cost of sex scale | LogNormal($\mu = -0.07, \sigma = 0.09$) | Gibson et al. (2017) |
| $s$ | Proportion of male offspring produced | 0.5 | Assumption |
| $b_U$ | Number of offspring produced by an uninfected host | 20 | Lively (2010b) |
| $b_I$ | Number of offspring produced by an infected host | $(1 - V)b_U$ | Lively (2010b) |
| $a_U$ | Density dependent effect coefficient of uninfected hosts | 0.001 | Lively (2010b) |
| $a_U$ | Density dependent effect coefficient of infected hosts | 0.001 | Lively (2010b) |
| $r_{\text{host}}$ | Host recombination probability | 0.2 | Lively (2010b) |
| $r_{\text{parasite}}$ | Parasite mutation probability | 0.05 | Assumption |
| $p_{\text{host}}$ | Probability that at least one sexual and asexual host enters the population | 0.1 | Assumption |
| $p_{\text{infected}}$ | Probability that at least one infected host enters the population | 0.02 | Assumption |

We use the Population Monte Carlo approach (Turner and Van Zandt, 2012), which allows for efficient sampling while ensuring that final result still converges to a correct (approximate) Bayesian posterior. The Population Monte Carlo approach begins with a basic ABC. For each random parameter sample drawn from the prior distribution, the model is simulated and a sample of a subpopulation is drawn from the simulated system, equal to the number of sites collected in a study. Summary statistics are calculated based on the last 100 generations out of 1100 generations; the current parameter set is accepted if the distance between simulated and observed data is less than a specified tolerance value. Distance is measured by the sum of absolute differences in summary statistics between simulated and observed data. This process is repeated until 100 parameter sets are accepted.

After the first run (*t* = 1), equal weights (*w*_*i*,1_ = 1/100) are assigned to each accepted parameter set ***θ***_*i*,1_ (*i* = 1, 2, …, 100). A weighted random sample (***θ***\*) is drawn from the accepted parameters of the previous run (*t* − 1) with weights *w*_*i,t*−1_. For any run *t* > 1, a parameter sample (***θ***_*i,t*_) is proposed from a multivariate normal distribution with a mean ***θ***\* and a variance covariance matrix that is equal to 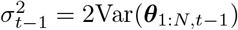, where Var(***θ***_1:*N,t*−1_) is the weighted variance covariance matrix of the accepted parameters from the previous run and *N* is the total number of accepted parameters from the previous run.

The parameter *G*_asex_ is rounded to the nearest integer and the model is simulated. If a proposed parameter is accepted, the following weight is assigned:

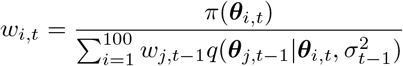

where *π*(·) is a prior density and 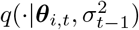 is a multivariate normal density with mean ***θ***_*i,t*_ and variance covariance matrix 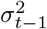. For each run, 100 parameters are accepted and weights are normalized at the end to sum to 1.

For each observed data set, we perform 4 runs with decreasing tolerance. First three tolerance values are chosen in a decreasing sequence to reach the final step quicker. The tolerance value of the final run is chosen so that a parameter set will be accepted if its each simulated summary statistic deviates from the corresponding observed summary statistic by 0.1 units on average. For spatial data (Dagan et al., 2013a; McKone et al., 2016), four summary statistics are compared: the mean proportion of infected and sexually reproducing hosts and CV in these proportions across populations. Tolerance values of 1.6, 0.8, 0.6 and 0.4 are used for each run. For spatiotemporal data (Vergara et al., 2014b), we additionally compare the CV of proportions of infected and sexually reproducing hosts across generations. In this case, larger tolerance values (2.4, 1.2, 0.9 and 0.6) are used for each run to account for a higher number of summary statistics being compared.

All statistical results are weighted by parameter weights of the final run; confidence intervals are obtained by taking weighted quantiles. However, it is always not possible to obtain the exact 95% confidence interval when the smallest (or the largest) value has a weight greater than 0.025. In these cases, we take the lowest (or largest) possible quantile that is closest to the 2.5% (or the 97.5%) quantile. While this procedure results in a slightly narrower confidence interval, the rounding errors in quantiles are negligible. This rule leads to one instance where we use the 2.82% quantile instead of the 2.5% quantile as the lower end of a confidence interval.

### 2.5 Power analysis

Using estimated parameters for each data set, we calculate the power to detect a significant positive correlation between infection prevalence and frequency of sexual hosts. For each parameter sample from the final run of the ABC, 10 simulations are run. For each simulation, we take the last two generations — assuming that a year contains two snail generations (estimated generation time is 4-9 months (Neiman et al., 2005)) — from the simulation and choose *n* populations at random from 40 simulated populations. For each selected population, hosts are divided into four categories based on their infection status (infected/uninfected) and reproductive mode (asexual/sexual), and the mean proportion of hosts in each category is calculated by averaging over two generations. Independent multinomial samples of size *m* are drawn from each selected population based on the proportions in every four categories. Correlation between the proportion of infected hosts and the proportion of sexual hosts is tested using the Spearman’s rank correlation at a 5% significance level.

While we do not explicitly try to match observed correlations, this procedure is essentially a retrospective power analysis. The main purpose of a power analysis is to design an experiment and determine an appropriate sample size (Cohen, 1992). If instead, power is calculated based on an observed effect size after an experiment, the estimated power is likely to be correlated with the observed significance of the result; therefore, interpreting significance (or a lack of significance) of a statistical result based on a retrospective power analysis is inappropriate (Goodman and Berlin, 1994; Senn, 2002).

Here, we do not try to justify the significant correlation observed by (McKone et al., 2016) or the nonsignificant correlation observed by (Dagan et al., 2013a) based on this power analysis. Instead, we use a power analysis to better understand how a study design or the dynamics of the model might affect the correlation between the prevalence of infection and sexual reproduction while using biologically realistic parameters.

## 3 Results

First, we compare observed summary statistics with either fitted or predicted summary statistics (Fig. 1). *Fitted* (filtered) summary statistics (Fig. 1) are values that have been accepted by the ABC procedure as being sufficiently close to observed summary statistics; these can be interpreted as model-based estimates of the summary statistics (and associated confidence intervals) of the study sites. *Predicted* (unfiltered) summary statistics, in contrast (Fig. 2), use parameters from the posterior distribution sampled by ABC, but generate new simulations of the dynamics and calculate summary statistics from randomly selected subpopulations. These values represent summary statistics sampled from the full distribution of coevolutionary dynamics that are consistent with the observed dynamics; they represent the expected range of dynamics from randomly sampled host-parasite metapopulations with a given distribution of parameters. Because they include an additional level of dynamical variability, the predicted summary statistics range much more widely than the fitted summary statistics.

**Figure 1:**
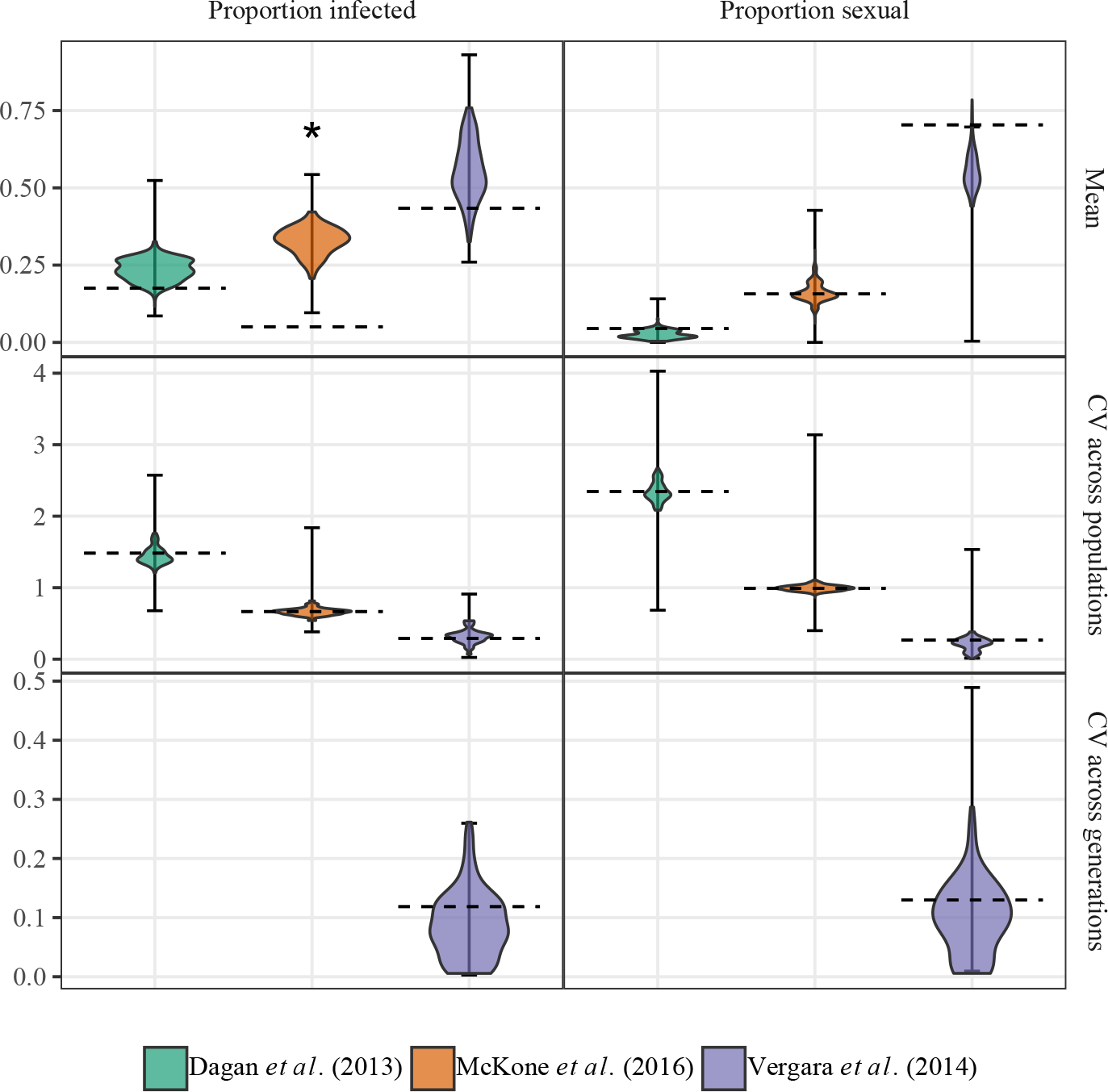
Summary statistics of the observed data vs. distribution of summary statistics of the simulated data from the posterior samples. Dotted horizontal lines represent observed summary statistics. Violin plots show the weighted distribution of fitted summary statistics (i.e., summary statistics that were accepted during ABC). Error bars show 95% weighted quantiles of predicted summary statistics. The weights correspond to the posterior distribution weights from ABC. For each posterior sample, 10 simulations are run and each simulated system is sampled at random 100 times so that each sample consists of the same number of populations as in the fitted data. All univariate summary statistics are matched reasonably well, except for the mean proportion of infected hosts in McKone et al. (2016) (indicated by an asterisk).

**Figure 2:**
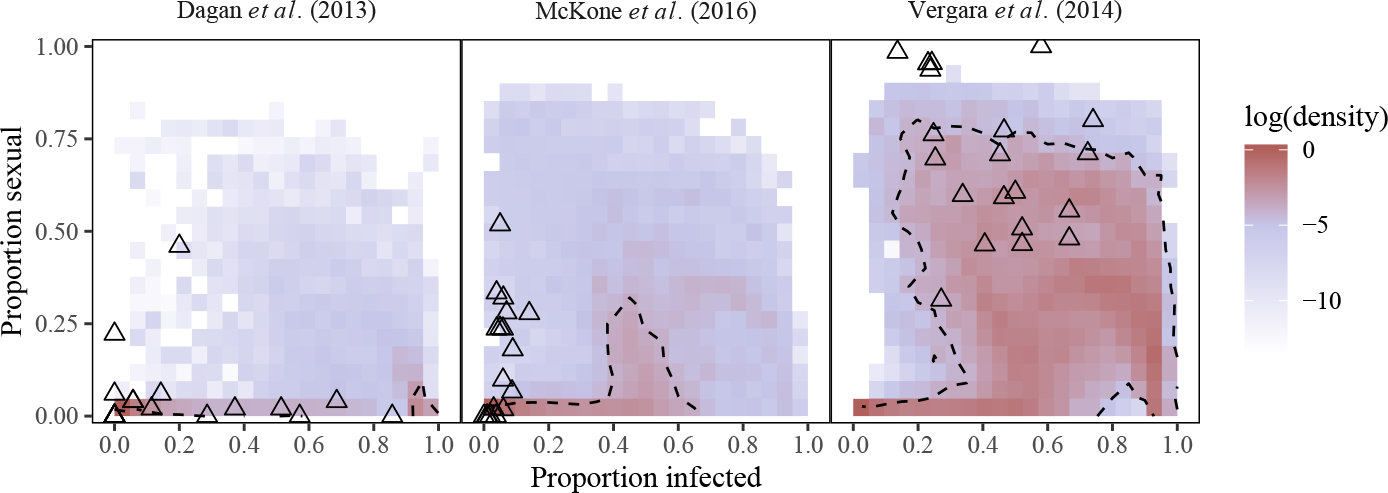
Predicted relationship between mean infection prevalence and mean proportion of sexual hosts in each population. For each posterior sample, 10 simulations are run. For each population within a simulation, mean infection prevalence and mean proportion of sexual hosts is calculated by averaging across last two generations. Each point, consisting of mean infection prevalence and meal proportion of sexual hosts, is assigned the weight of the parameter used to simulated the population. The density at each cell is calculated as the sum of weights of the points that lie within it. Cell densities are normalized by dividing by the maximum grid densities for each fit. Dashed contours are calculated at log(density) = −4. Open triangles represent observed data; proportion of sexual hosts is computed by doubling proportion of male hosts.

Our simple meta-population Red Queen model can capture observed variation in infection prevalence and frequency of sexual hosts reasonably well; both temporal and spatial variation (measured by CV across mean proportions) are well-matched by the model. However, as model fitting is performed by minimizing the sum of absolute distance between observed and simulated summary statistics, our method does not guarantee that all summary statistics are equally well-fitted. The model tends to overestimate the mean proportion of infected hosts. The observed mean proportions of infected hosts are 0.175 (Dagan et al., 2013a), 0.051 (McKone et al., 2016), and 0.440 (Vergara et al., 2014b), whereas their ABC-based estimates are 0.240 (95% CI: 0.174 - 0.286), 0.313 (95% CI: 0.233 - 0.404), and 0.542 (95% CI: 0.360 - 0.730), respectively. The model underestimates the mean proportion of sexual hosts for Dagan et al. (2013a) and Vergara et al. (2014b). Observed mean proportions of sexual hosts are 0.045 and 0.704, respectively, whereas their ABC-based estimates are 0.026 (95% CI: 0.007 - 0.048) and 0.596 (95% CI: 0.441 - 0.679).

To further diagnose the fit, we compare the predicted relationships between the mean proportion of infected hosts and the mean proportion of sexual hosts across subpopulations with the observed data (Fig. 2). Despite its accuracy in reproducing summary statistics reported by Dagan et al. (2013a), our model poorly captures the relationship between the mean proportion of infected hosts and the mean proportion of sexual hosts (Fig. 2; Dagan et al. (2013a)). The model predicts sexual reproduction to be well maintained when infection prevalence is high (> 40%) whereas the observed data (Dagan et al., 2013a) suggests that sexual reproduction can only be supported when infection prevalence is low (< 20%). McKone et al. (2016) also found sexually reproducing snails in sites with low infection prevalence (< 20%).

On the other hand, Vergara et al.’s (2014b) data set suggests that sexually reproducing populations are likely to have a relatively high prevalence of infection (approximately 20% - 80%). This observation is broadly consistent with our model prediction. Most of the data points fall within the range of model predictions (Fig. 2; Vergara et al. (2014b)). However, there is one site in which more than 90% of the snails were found to be sexual throughout the study period of 5 years (Vergara et al., 2014b); our model fails to maintain such high levels of sexual reproduction.

There is a high posterior density region in which the proportion of infected hosts remains almost constant (around 0.5) among subpopulations while the proportion of sexual hosts can range from 0 to 0.3 (visible in the fits to McKone et al. (2016)). As transmission rate (*β*) increases, selection for sexual hosts increases but the increasing number of resistant offspring prevents further infection from occurring and can decrease overall infection prevalence. This trend is consistent with previous results of Lively (2001) who noted that there is a region in which either sexual and asexual reproduction can be selected exclusively under the same infection prevalence.

The proportion of sexual hosts decreases when infection prevalence is extremely high (visible in the fits to Vergara et al. (2014b)). This pattern can be explained by the decrease in fitness of sexual hosts with the increase in infection prevalence, predicted by Ashby and King (2015). The same pattern can be seen in earlier work by Lively (2010b) although it was not discussed.

Fig. 3 presents parameter estimates. Keeping in mind that we do not obtain good fits to data from Dagan et al. (2013a) and McKone et al. (2016), we still find that high virulence and a low ratio of asexual to sexual genetic diversity are necessary to explain the observed dynamics. Moreover, we are able to capture observed differences in mean and variation in infection prevalence among studies in our estimates of transmission rate parameters (*β*_mean_ and *β*_CV_).

**Figure 3:**
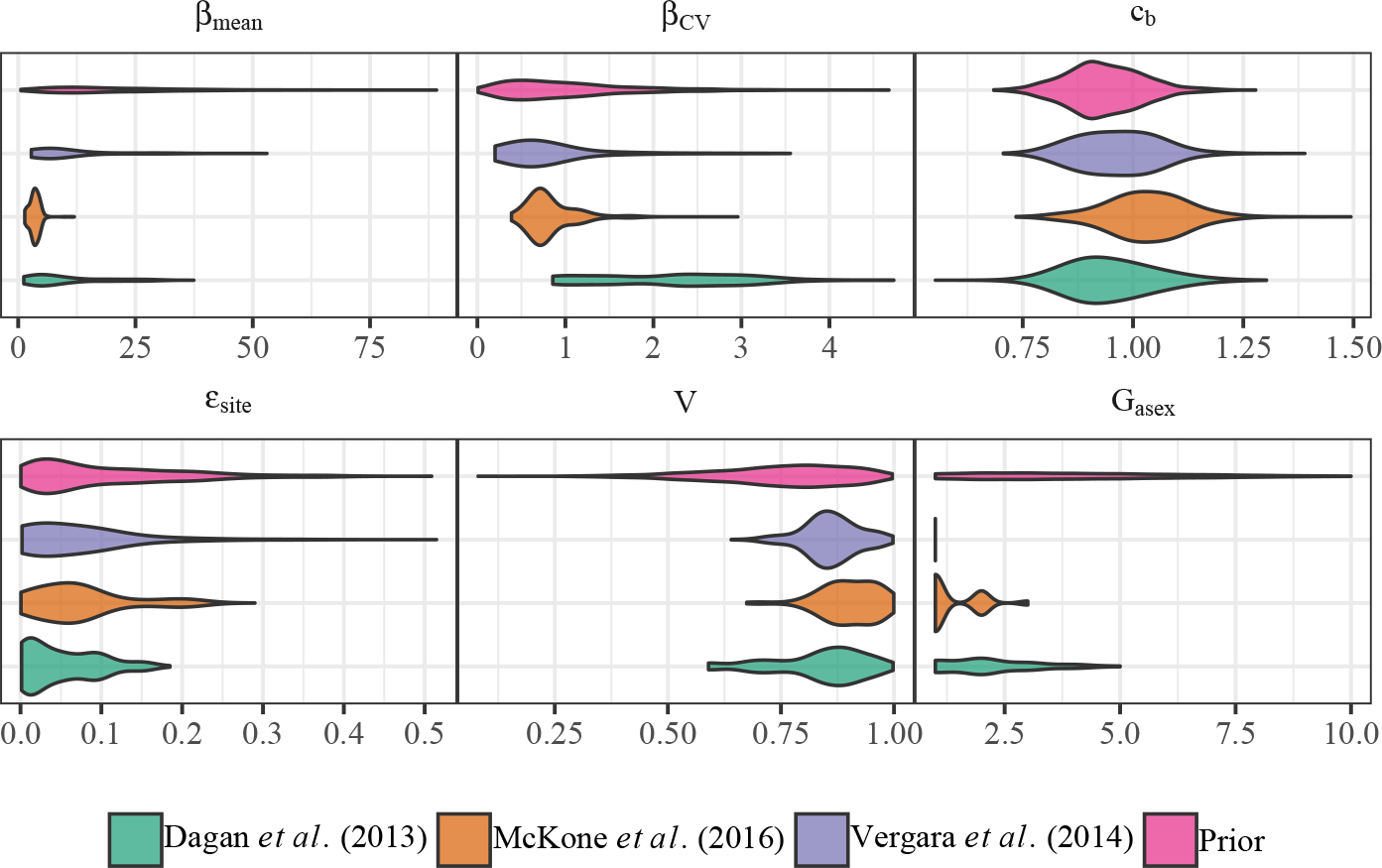
Parameter estimates from Sequential Monte Carlo Approximate Bayesian Computation. We estimate mean transmission rate (*β*_mean_), CV in transmission rate (*β*_CV_), virulence (*V*), mixing proportion (*ϵ*_mean_), number of asexual genotypes (*G*_asex_), and a scale parameter for cost of sex (*c*_*b*_). Violin plots represent weighted distribution of 100 posterior samples obtained from ABC. Violin plots for prior distribution is obtained by drawing 10000 random parameter samples from the prior distribution. *G*_asex_ is a discrete variable but is drawn on a continuous scale for convenience. See Table 1 for full parameter descriptions and their prior distributions.

Our fits to McKone et al. (2016) suggest that the scale parameter for the cost of sex, *c*_*b*_, should be higher than our prior assumption based on Gibson et al. (2017) that estimated the cost of sex to be 2.14 (95% CI: 1.81 - 2.55). Ashby and King (2015) defined *c*_*b*_ as additional costs and benefits of sex, where *c*_*b*_ = 1 corresponds the two fold cost. Under their interpretation, our estimate of *c*_*b*_ corresponds to a slightly lower estimate of the cost of sex: 1.95 (95% CI: 1.68 - 2.4). (We propose an alternate interpretation to this parameter estimate below).

Finally, our power analyses suggest that there is high power to detect a positive correlation between infection prevalence and frequency of sexual hosts in the systems studied by Dagan et al. (2013a) and McKone et al. (2016) (Fig. 4). Such high power predicted for Dagan et al. (2013a) is particularly surprising given that they were not able to observe the expected correlation. This discrepancy implies that the snail populations studied by Dagan et al. (2013a) show sufficient variation in infection prevalence in order for the correlation to be observed under pure Red queen selection, but other underlying factors that are neglected by our model may have caused the populations to deviate from their expected behaviors. On the other hand, our model predicts low power for detecting the positive correlation for the system studied by Vergara et al. (2014b) (Fig. 4).

**Figure 4:**
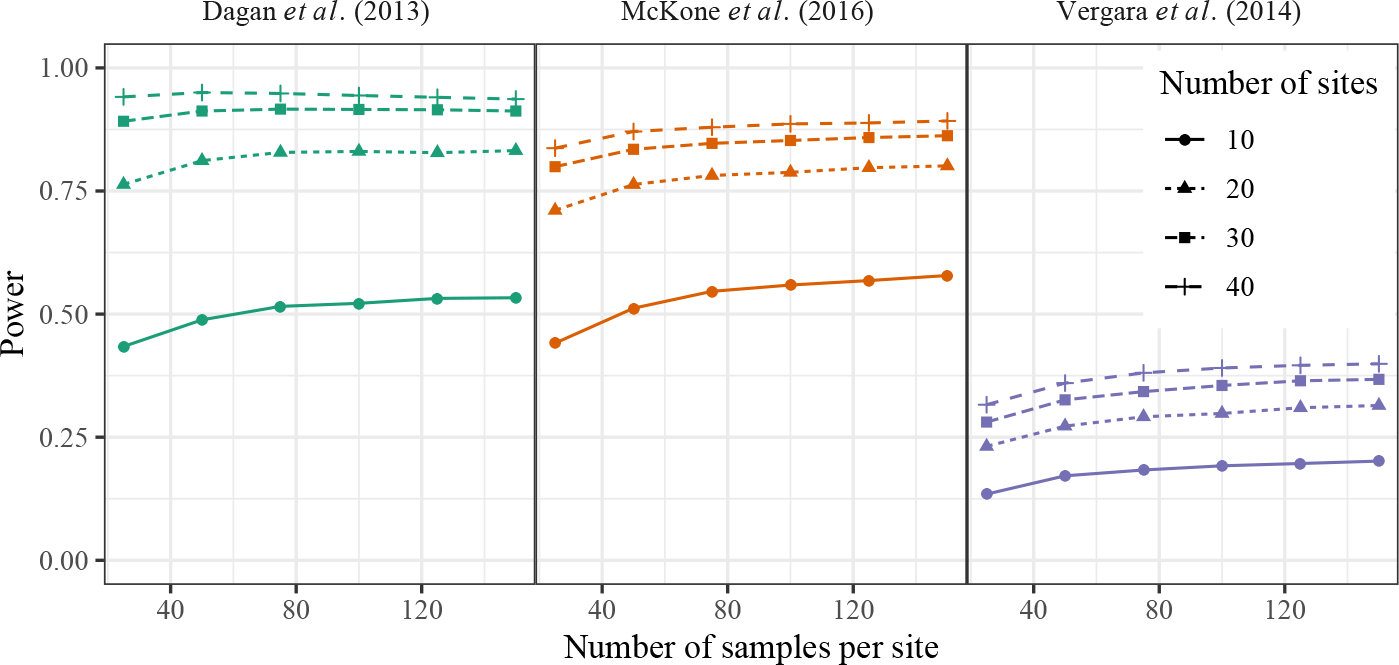
Power to detect a statistically significant positive correlation between infection prevalence and frequency of sexual hosts. Spearman’s rank correlation was used to test for correlation between infection prevalence and frequency of sexual hosts in simulated data from the posterior distributions.

Overall, our analysis suggests that increasing the number of study sites is a more effective way to increase power than increasing the number of samples per site (Fig. 4). The correlation between the proportion of infected and sexual hosts depends on the spatial distribution of parasite infection; increasing the number of sites can better capture this distribution and increase the power to observe the correlation. On the other hand, collecting more samples from a site only gives us a more accurate estimate of the two proportions for that specific site; it does not tell us how these proportions are distributed across space. As a result, the power quickly saturates once we have a sufficient number of samples (> 100) to estimate the two proportions reliably at each site.

While we originally planned to perform power analysis using Spearman’s rank correlation, we repeated the analysis using Pearson’s correlation after applying arcsine square root transformation (Lively, 1992) to see whether this procedure improves power. Using Pearson correlation gives slightly higher power to detect the positive correlation between frequency of sexual hosts and infection prevalence (see Appendix).

## 4 Discussion

A simple metapopulation model for host-parasite coevolution is able to match the observed prevalence of sexual reproduction and trematode infection in snail populations (Fig. 1). A direct comparison between a model and a data set allows us to infer biologically meaningful parameters of a model (Fig. 3) and estimate the power to observe a significant, positive correlation between the proportion of infected hosts and the proportion of sexual hosts (Fig. 4), as predicted by Lively (1992). However, discrepancies between the model predictions and observed data suggest that a simple host-parasite coevolution model may not be able to sufficiently explain the maintenance of sexual reproduction observed in snail populations (Fig. 2).

A model that fits poorly can sometimes tell us more about a biological system than a model that fits well. Our model clearly failed to match the data presented by Dagan et al. (2013a) (Fig. 2). The snail populations studied by Dagan et al. (2013a) live in qualitatively different environments from the two other snail populations that we considered (Vergara et al., 2014b; McKone et al., 2016). For example, their habitats are subject to seasonal flash floods (Ben-Ami and Heller, 2007), which can affect reproductive strategies of snails and interfere with the host-parasite coevolution (Lytle, 2000; Ben-Ami and Heller, 2007). As a result, a positive correlation between infection prevalence and frequency of sexual reproduction could not be detected from the system even though our model (which does not incorporate disturbance) predicts high power. These results suggest that a significant correlation (or the lack of it) between infection prevalence and the frequency of sexual reproduction may not provide sufficient evidence for (or against) the role of parasites in maintaining sexual reproduction in the host population.

The model fits to Dagan et al. (2013a) and McKone et al. (2016) suggest that the cost of sex can be overcome and sexual reproduction can be maintained only if infection prevalence is much higher than the observed prevalence (Fig. 2). In other words, the benefit of producing offspring with novel genotype is relatively small when infection prevalence is low. In order to support sexual reproduction at lower infection prevalence, the benefit of sex must be greater or other mechanisms must compensate for the difference. As our model relies on a simple structure and strong parametric assumptions, an additional benefit of sex can only be provided by lowering the cost of sex (i.e., increasing the scale parameter, *c*_*b*_).

The simple structure of the model and limited genetic diversity can explain the discrepancy between model prediction and the observed data by McKone et al. (2016). Here, we assumed that host resistance to infection is determined entirely by two biallelic loci, resulting in 10 possible genotypes. However, it is unlikely that such a simple model can capture the genetic interaction between hosts and parasites observed in nature. Although exact genetic architecture that determines trematode infection in snails (e.g., the number and inheritance of loci involved in parasite resistance) is not known, the documented genetic diversity of snails is far greater than what our model assumes (Fox et al., 1996; King et al., 2011; Dagan et al., 2013a). Increasing the maximum possible genetic diversity of the model would have allowed sexual hosts to escape infection more easily and maintained sexual reproduction at a lower prevalence of infection (Lively, 2010a; King and Lively, 2012; Ashby and King, 2015).

While the positive correlation between frequency of sexual hosts and prevalence of infection can provide evidence for the effect of the parasite on the maintenance of sexual reproduction in the host population, it does not fully capture the coevolutionary process. In particular, the positive correlation represents a contrast between populations that undergo Red Queen dynamics and those that do not (rather than a continuous relationship between the frequency of sexual hosts and prevalence of infection): low-risk populations will be dominated by asexual individuals whereas high-risk populations will consist of both sexual and asexual individuals. Lively (2001) gave a similar explanation and predicted that large variation in the risk of infection is required to observe the predicted correlation.

The similarity between the correlations predicted by the model and the correlations observed in nature may provide us more confidence that the observed populations provide evidence for the Red Queen Hypothesis (Fig. 5). However, the strength of the correlation is not a direct measure of the strength of selection imposed on the host population. Moreover, when populations are actively coevolving, wide range of correlations can be detected (Fig. 5: Vergara et al. (2014b)). While simple statistical summaries, such as a correlation coefficient, are easily accessible, more sophisticated and mechanistic statistical models that yield biologically interpretable parameters may be better suited for understanding the effect of parasites on the maintenance of sexual reproduction in the host population.

**Figure 5:**
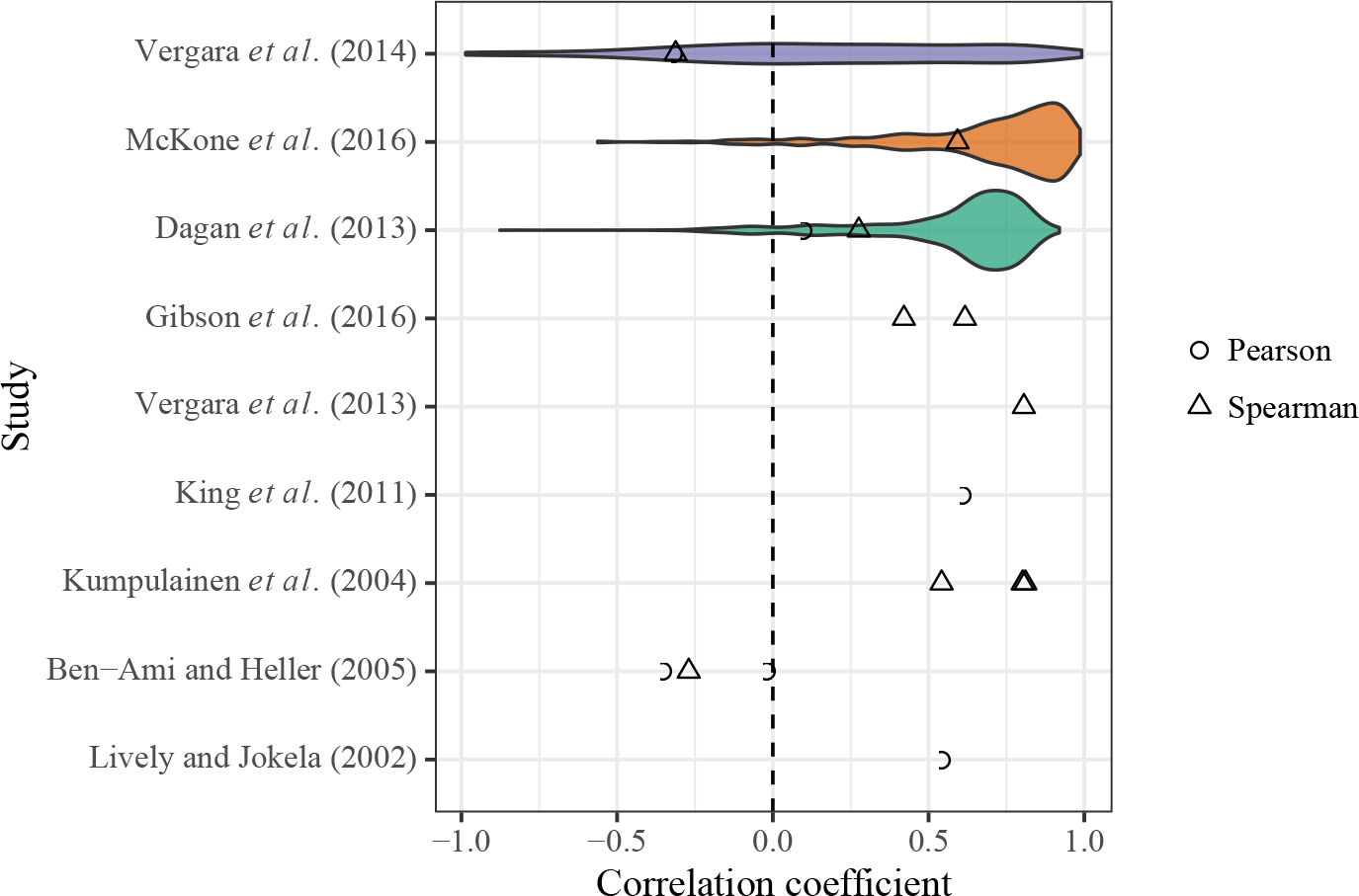
Predicted vs. observed correlation between infection prevalence and frequency of sexual hosts. Violin plots show weighted distribution of predicted strength of correlation based on ABC fits. Predicted correlation was measured for each simulation from the posterior by taking into account the last two generations from each simulated population. Triangles and circles represent observed correlation sizes from previous studies. Dagan et al. (2013a) and Vergara et al. (2014b) do not report the size of the correlation; correlations are re-calculated from their data sets using both Pearson (after applying arcsine square root transformation) and Spearman correlations.

Mathematical models have used extensively to build theoretical foundations for the evolution of sex but only a few models have been confronted with data. The idea of the two-fold cost of sex was only recently tested directly by fitting a theoretical model to observed data (Gibson et al., 2017). Making statistical inference on the observed systems and testing theoretical models against data may provide deeper insight into underpinnings of maintenance of sexual reproduction.

## 5 Acknowledgements

We thank SHARCnet for providing computational resources.

## 6 Funding

This work is supported by The Natural Sciences and Engineering Research Council, Undergraduate Student Research Award (to SWP).

## Appendix

**Figure A1:**
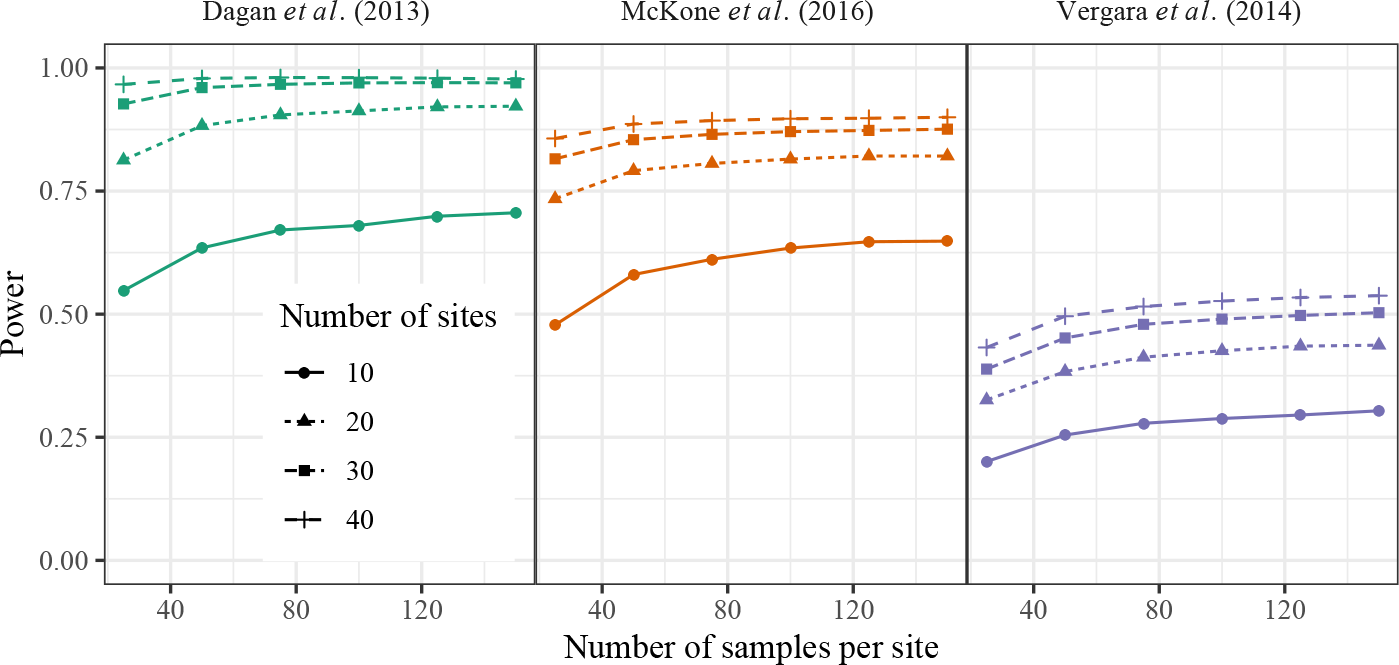
Power to detect a statistically significant positive correlation between infection prevalence and frequency of sexual hosts. Pearson correlation was used to test for correlation between square root arcsine transformed infection prevalence and frequency of sexual hosts in simulated data from the posterior distributions.

